# Structural basis of human PRPS2 filaments

**DOI:** 10.1101/2022.07.11.499506

**Authors:** Guangming Lu, Huan-Huan Hu, Chia-Chun Chang, Jiale Zhong, Xian Zhou, Chen-Jun Guo, Tianyi Zhang, Yi-Lan Li, Boqi Yin, Ji-Long Liu

## Abstract

PRPP synthase (PRPS) transfers the pyrophosphate groups from ATP to ribose-5-phosphate to produce 5-phosphate ribose-1-pyrophosphate (PRPP), a key intermediate in the biosynthesis of several metabolites including nucleotides, dinucleotides and some amino acids. There are three PRPS isoforms encoded in human genome. While hPRPS1 and hPRPS2 are expressed in most tissues, hPRPS3 is exclusively expressed in testis. Although hPRPS1 and hPRPS2 share 95% sequence identity, hPRPS2 has been shown to be less sensitive to allosteric inhibition and specifically upregulated in certain cancers in the translational level. Recent studies demonstrate that PRPS can form a subcellular compartment termed the cytoophidium in multiple organisms across prokaryotes and eukaryotes. Forming the cytoophidium is considered as a distinctive mechanism involving the polymerization of the protein. In order to investigate the function and molecular mechanism of hPRPS2 polymerization, we solve the polymer structure of hPRPS2 at 3.08Å resolution using cryo-Electron Microscopy (cryo-EM). hPRPS2 hexamers stack into polymers in the conditions with the allosteric/competitive inhibitor ADP. The binding modes of ADP at the canonical allosteric site and at the catalytic active site are clearly determined. A point mutation disrupting the inter-hexamer interaction prevents hPRPS2 polymerization and results in significantly reduced catalytic activity. Our findings suggest that the regulation of hPRPS2 polymer is distinct from *E. coli* PRPS polymer and provide new insights to the regulation of hPRPS2 with structural basis.

## INTRODUCTION

PRPP synthase (PRPS) transfers the pyrophosphate groups from ATP to ribose-5-phosphate to produce 5-phosphate ribose-1-pyrophosphate (PRPP). PRPP is important for de novo purine and pyrimidine nucleotide metabolism and salvage pathway (Li et al., 2015). PRPP is also used in the biosynthesis of amino acids histidine and tryptophan, NAD and NADP (Hove-Jensen, 1988; 1989). Moreover, PRPP is also used in the biosynthesis of methotrexate in archaea and pentose polyphenylphosphate in *Mycobacterium tuberculosis* (Graham and White, 2002; Wolucka, 2008). PRPS is very conservative in evolution and widely exists in all three life domains, such as bacteria, archaea and eukaryotes. PRPS can be divided into several types in different species.

Human PRPS (hPRPS) belongs to the first type of PRPS, which can only use ATP or dATP as pyrophosphate donor, and can be allosterically regulated by inorganic phosphate Pi and ADP (Hove-Jensen and McGuire, 2004). There are three genes encoding PRPS in human body, *prps1, prps2, prps3*, which encode hPRPS isoenzymes 1-3 respectively. Human *prps1* and *prps2* are located on the X chromosome and are expressed in all tissues. Human *prps3* is located on chromosome 7 and is only expressed in testis (Taira et al., 1989).

There are two subtypes of hPRPS2 in vivo. hPRPS2-long has three more amino acids than hPRPS2-short after the 102_nd_ amino acid. Both hRRPS1 and hPRPS2-short have 318 amino acids, with a similarity of 95%. Although hRRPS1 and hPRPS2 are highly similar, they still have some different properties. For example, hPRPS2 is more sensitive to heat inactivation, the saturation concentration of substrate is higher than that of hPRPS1, and it is less likely to be inhibited by nucleoside diphosphate acid (Nosal et al., 1993).

In human body, hPRPS1 and hPRPS2 have different functions. The activity stability of hPRPS1 is very important to the body. When the mutation activity of hPRPS1 increases, it will cause hyperuricemia, myasthenia, gouty arthritis or neurosensory defects. When the mutation activity of hPRPS1 decreases, it will lead to neuropathy, deafness or intellectual disability.

Compared with hPRPS1, hPRPS2 plays a more important role in the occurrence and maintenance of cancer caused by the transcription factor Myc. The increase of nucleotide synthesis will inhibit hPRPS1, while the feedback inhibition of hPRPS2 on nucleotides is less obvious. Moreover, *prps2* gene has one more pyrimidine rich translation element (prte) in the 5 ’- noncoding region than *prps1* gene. Eukaryotic initiation factor (eIF4E) and other possible factors interact with prte to increase the mRNA encoding *prps2*, thereby increasing the concentration of hPRPS2 to increase the synthesis of nucleotide. These evidences show that hPRPS2 plays an important role in the metabolism of cancer cells with high expression of myc (Cunningham et al., 2014).

There is a crystal structure of hPRPS1 (Li et al., 2007), but there is no crystal structure of PRPS2 at present. Using light microscopy, Wilhelm et al. found that PRPS formed filamentous structures in a variety of eukaryotic cells such as yeast, *Drosophila* oocytes, rat neurons, human fibroblasts and zebrafish retinal epithelial cells (Begovich et al., 2020; Noree et al., 2019). Recently, we also found that PRPS can form filamentous structures in prokaryotes such as *Escherichia coli* in vitro and in vivo (Hu et al., 2022). We further understood the structure *E. coli* PRPS at near atomic resolution through cryo-EM.

It is not clear whether hPRPS2 can form filaments. Here, we obtain the structure of hPRPS2 at 3.08A resolution using cryo-EM. In this structure, we can see a clear hexamer, and we have also found the key amino acid residues at the interface of adjacent hexamers. Point mutations at the interface can destroy filament assembly, resulting in decreased enzyme activity. This result supports the views we previously observed in prokaryotic cells (Hu *et al*., 2022).

In conclusion, our work reveals the structure of hPRPS2 for the first time and finds that it can form filaments. Comparing the PRPS structure of *E. coli* obtained before, we find that hPRPS filament was similar to *E. coli* type A filament. Although hPRPS1 and hPRPS2 have 95% identical amino acid residues, structural comparison shows that 5% of the different amino acid residues locate on the surface of the hexamers. In addition, we analyze the active sites and allosteric regulatory site of hPRPS2 filaments. Our results support that filamentation of metabolic enzymes provides an additional layer of regulation for metabolism.

## RESULTS

### hPRPS2 assembles into filaments

The filament structures of *E. coli* PRPS have recently been solved. However, the structure of hPRPS2 and whether it can form filament structure are still unclear. We purified the short isoform of hPRPS2 and tried different conditions to induce polymerization in vitro. Unlike *E. coli* PRPS, hPRPS2 only form filaments when incubated with ADP (2 mM) and Mg^2+^ (10 mM) (**Supplementary Figure S1**).

Using cryo-EM and single particle analysis, we solved the filament structure of hPRPS2 with the central layer map was estimated to be 3.08 Å (**Figure 1A; Supplementary Figure S2**). In the reconstructed model, hPRPS2 also forms a hexamer structure, such as hPRPS1 or other class I PRPS (**Supplementary Figure S3**). Hexamer is the basic unit of filament polymerization (**Figure 1B**). The twist and rise of hPRPS2 filament are 30° (left-handed twist) and 63 Å, respectively (**Figure 1C**).

**Figure 1.**
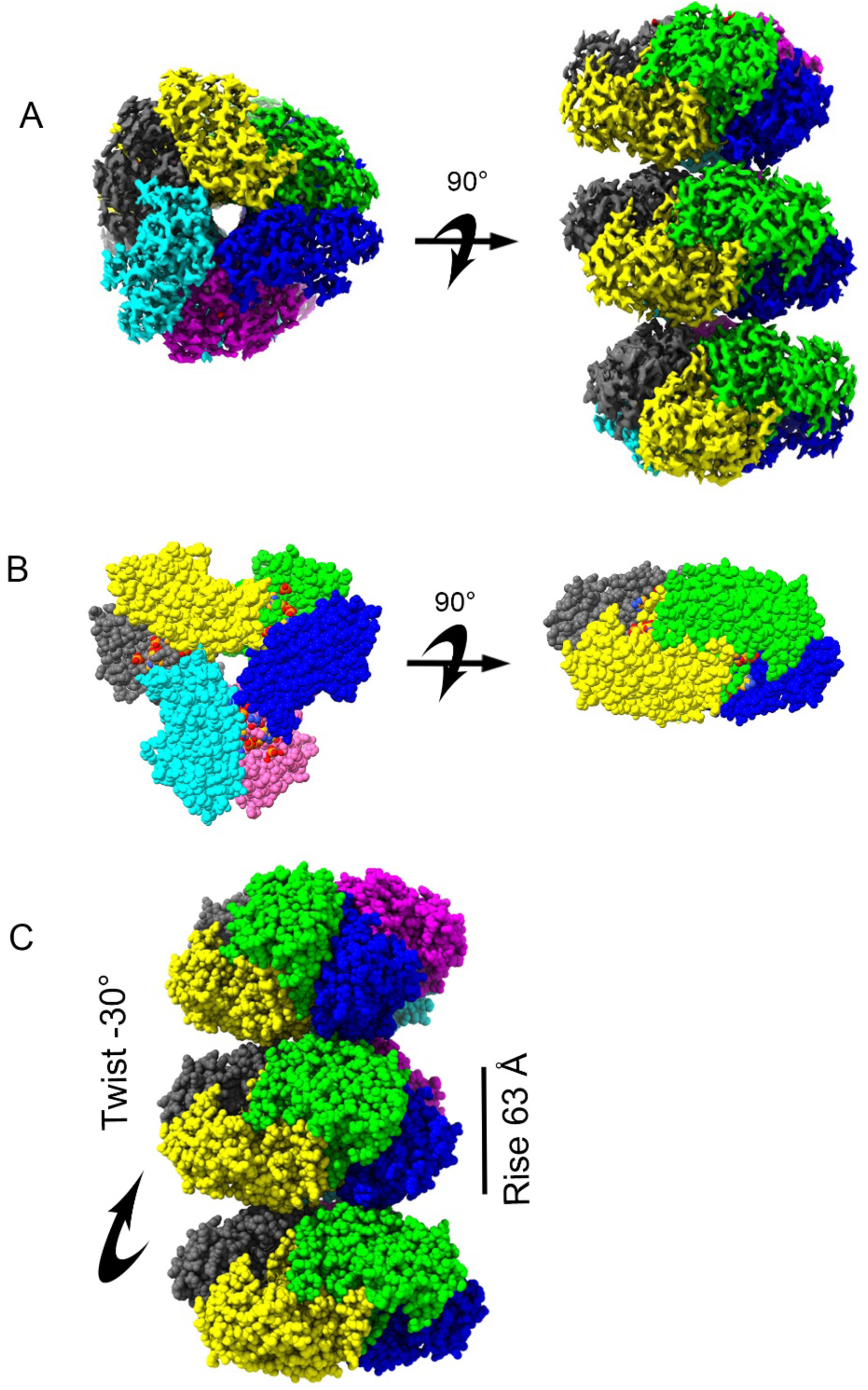
Overall structure of human PRPS2 filament. (A) The electron density map of type A filament (3.08 Å resolution). (B) Cryo-EM reconstruction of human PRPS2 hexamer. The hexamer is the unit of filament. Each chain is in different color. (C) The reconstruction structure of human PRPS2 filament. The rise of human PRPS2 filament is 63 Å. When hexamers are aggregated into filament, the adjacent hexamer is twisted by 30°.

### Ligand binding modes in hPRPS2 filaments

In the hPRPS2 filament, ADP was found at ATP binding site and allosteric site 1. There is a Pi occupied at the phosphate binding region of R5P, which may come from the hydrolysis of ADP or be preserved during protein purification (**Figure 2A and B**). Pi at the R5P active site forms hydrogen bonds with T225, T228 and the backbone of G227. ADP at the ATP active site is associated with an Mg^2+^, similar to that in other class I PRPS proteins. ADP in chain D forms hydrogen bonds with N37 and E39 in chain C, and there is π- π interaction between F35 in chain C and adenine base. R99 and H131 form salt bridges with the β-phosphate and α-phosphate separately. K176 in chain E also forms salt bridge with β-phosphate (**Figure 2C**).

**Figure 2.**
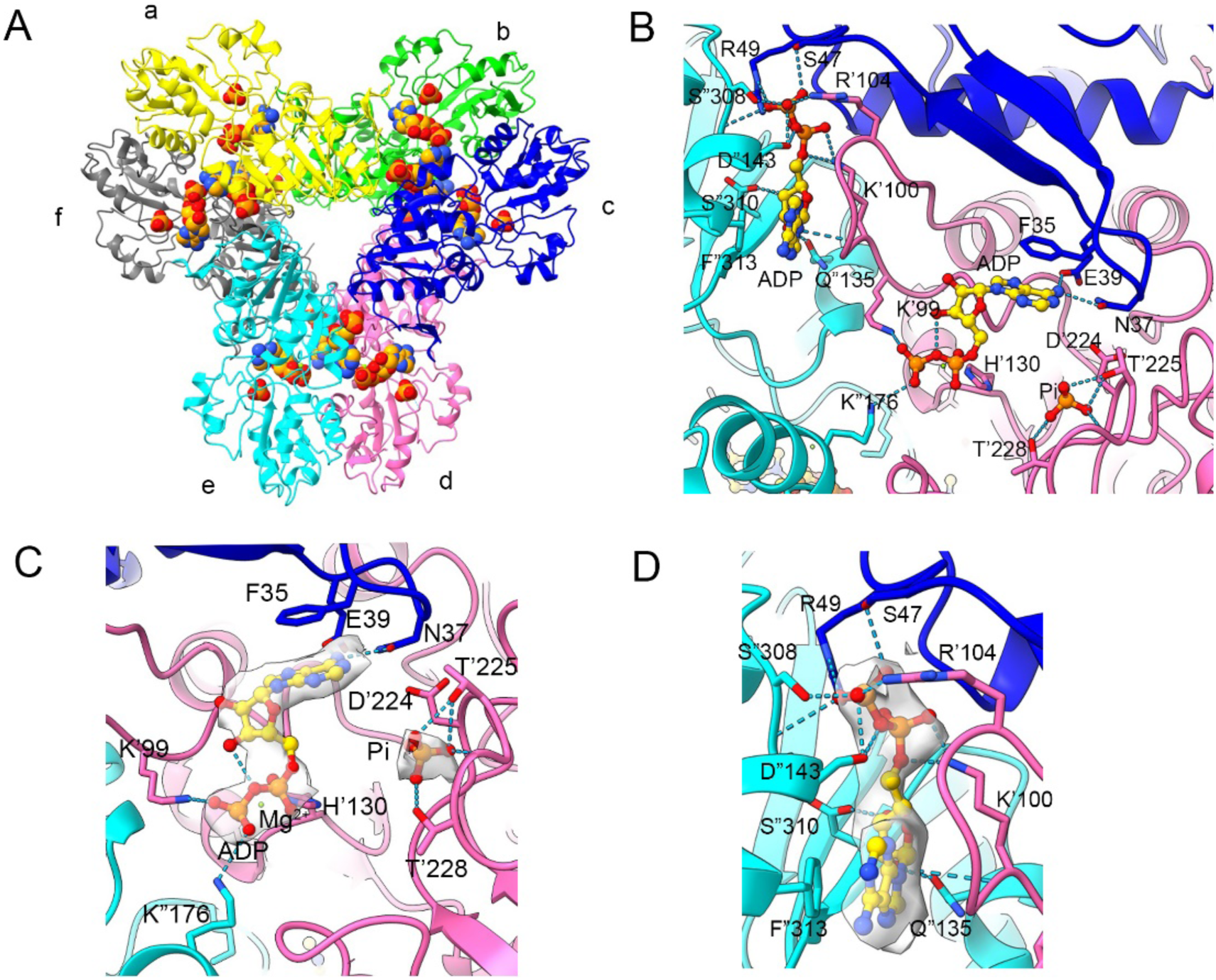
Ligand binding modes in human PRPS2 filaments. (A) Hexamer of human PRPS2 filaments. Each chain is marked with a different color. (B) ADP is identified on the allosteric site and active site of human PRPS2 filaments, while R5P binding site is bound by Pi. The residues that interact with ligands are indicated. Residues in chain d number with the ‘ symbol and in chain e number with the “ symbol. (C) Ligands of the active site of human PRPS2 filaments. ADP and Mg^2+^ occupy ATP binding sites at active sites. Pi can also be seen in the active site. Each chain is marked with a different color. (D) ADP in allosteric site of human PRPS2 filaments. Each chain is marked with a different color.

ADP in chain E allosteric site forms hydrogen bond with S47 in chain C, and S308, S310 in chain E through its β-phosphate. Q135, the backbone of K100 and D143 in chain E also form hydrogen bonds with the C-1 and C-2 hydroxyl groups respectively. In addition, there is also π-π interaction between F313 of chain E and adenine base. Salt bridges are formed between R49 in chain C, R104 in chain D with β-phosphate, and K100 in chain D with α-phosphate (**Figure 2D**).

### Contacts of hexamers in hPRPS2 filaments

The filament of hPRPS2 is stacked by hexamers with D3 symmetry. There are three interaction sites between two adjacent hexamers. Each interaction site contains some of the same amino acids (**Figure 3A**). hRPPS2 hexamers are connected by salt bridges formed between R301 and E298 pairs, the hydrogen bonds between R301, N305, and E307, and also the van der Waals’ Forces between R302 and R301 (**Figure 3B**).

**Figure 3.**
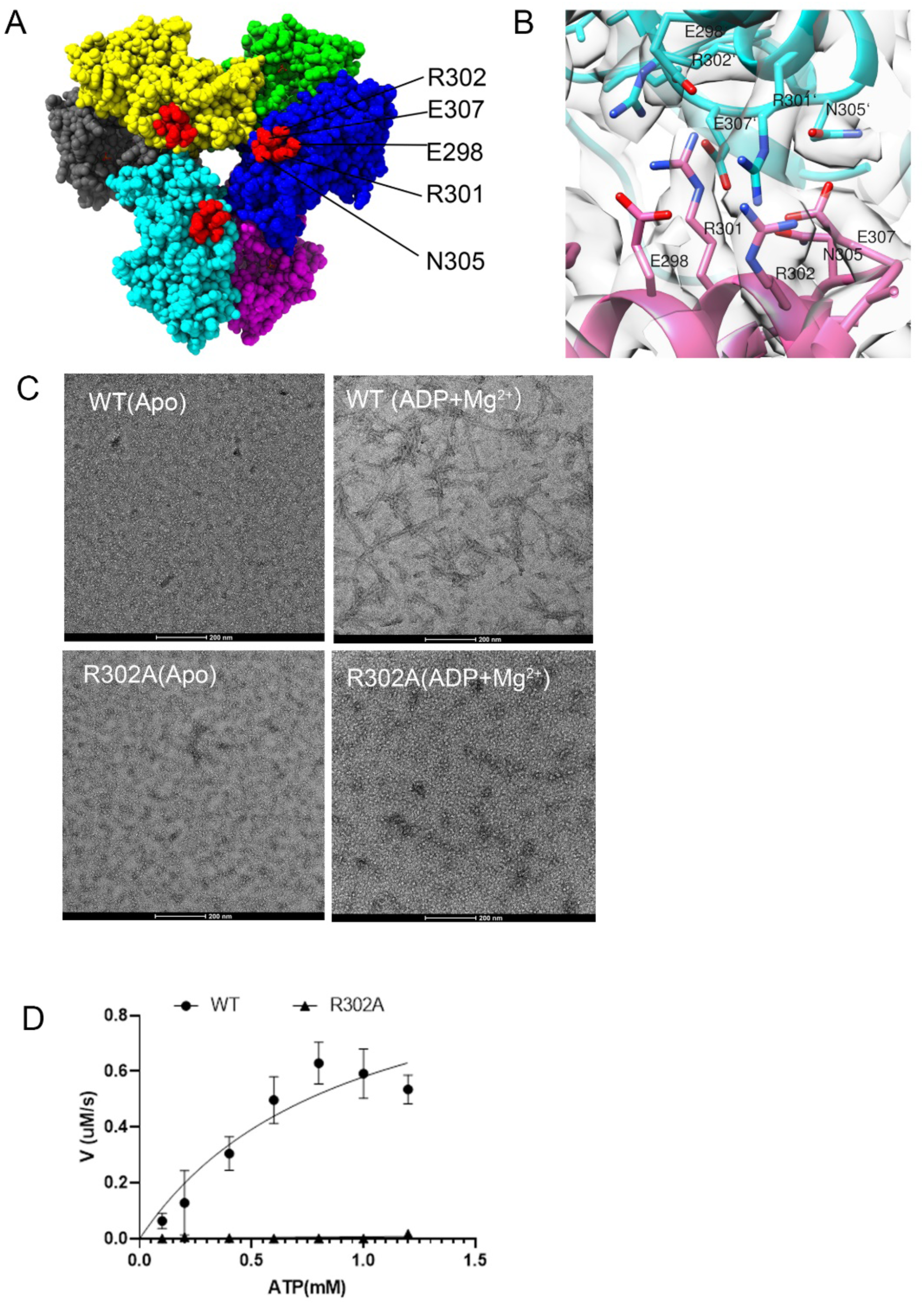
Contacts of hexamers in human PRPS2 filaments. (A and B) Maps and models of human PRPS2 filaments between adjacent hexamers. Residues responsible for the interactions are indicated. (C) Negative staining of human PRPS filaments. Wild type human PRPS2 can form filament under condition of ADP and Mg^2+^. And mutant R302A can disrupt the formation of filament. (D) Enzyme activity assay of wild type human PRPS2 and mutant R302A. Mutant R302A almost lost its activity.

After we mutated the amino acids at the interface between two adjacent hexamers, we found that the mutated hPRPS2^R302A^ could disrupt the formation of filament (**Figure 3C**). The filament forming ability was evaluated by negative staining electron microscopy. When any ligand was added to the solution, neither wild-type hPRPS2 nor mutant hPRPS2^R302A^ could form filaments. When incubated with allosteric regulator ADP and magnesium, hPRPS2 could form long filaments, and the mutant hPRPS2^R302A^ lossed its filament-forming ability.

To investigate the function of hPRPS2 filament, we used the coupled reaction method to test enzyme activity in vitro. Phosphoribosyltransferase (OPRT) can consume PRPP in the reaction orotate (OA) + PRPP → orotidine 5’-monophosphate (OMP) + PPi. OA has absorption at 295 nm, and the production of PRPP can be measured by the consumption of OA. The enzyme activity of hPRPS2 depends on the activation Pi. When the concentration of Pi was less than 10 mM, hPRPS2 did not show any activity. When Pi concentration was higher than 30 mM, hPRPS2 catalyzed the reaction at maximum velocity (**Supplementary Figure S4**). Therefore, we added 30 mM Pi reaction mixture to find the suitable substrate concentration. Finally, we tested the enzyme activity using 1 mM ATP, 1 mM R5P and 30 mM Pi. Our enzyme activity results showed that the mutant hPRPS2^R302A^ almost lost its activity.

### Comparison of hPRPS1 and hPRPS2 hexamers

We compared the sequences of hPRPS1 and hPRPS2, which have only 15 amino acid residues differences. We labeled these different amino acid residues in the structure of hPRPS2 hexamer (**Figure 4A**). Almost all the different amino acid residues are located on the surface of the hexamer. They are far away from active sites, allosteric sites and hexamer-hexamer interaction sites.

**Figure 4.**
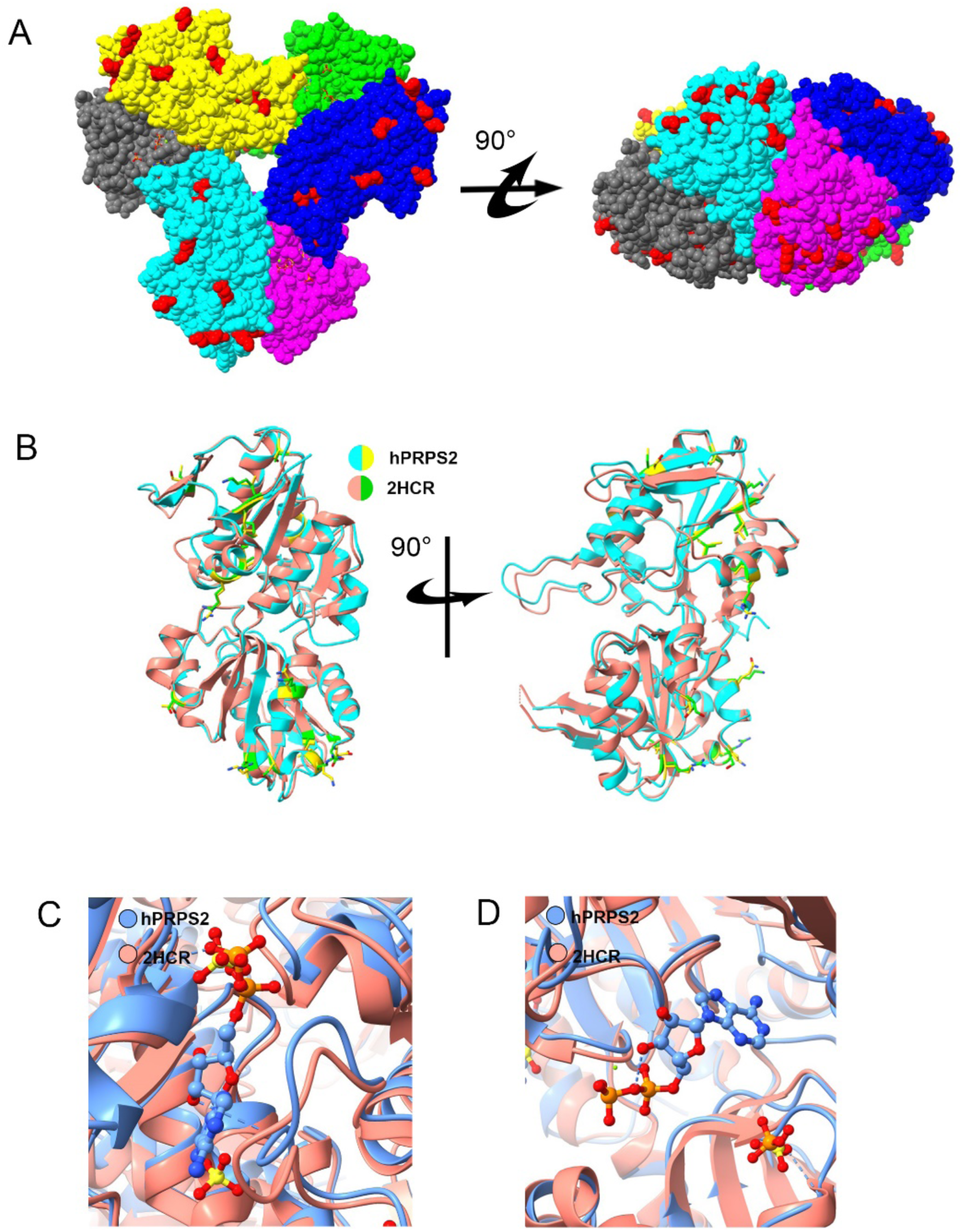
Structural comparison of hPRPS2 wth hPRPS1 (2HCR). (A) The difference of amino acids between hPRPS1 (2HCR) and hPRPS2. There are 15 amino acid differences between human PRPS1 monomer and human PRPS2 monomer. Most of the different amino acids are located on the surface of the hexamer. The amino acids with differences are marked in red. (B) Comparison of hPRPS1 and hPRPS2 monomer. The monomer of human PRPS1 is in red and human PRPS2 is in cyan. The amino acids with differences are green and yellow in human PRPS1 and human PRPS2, respectively. (C) Comparison of allosteric site and RF loop. There is an ADP at the allosteric site of human PRPS2, while SO_4_^2-^ in human PRPS1 can bind to allosteric site and another site. The chain of human PRPS1 (2HCR) is in red and of human PRPS2 is in blue. (D) Comparison of the active site between human PRPS1 (2HCR) and human PRPS2. In human PRPS2, ADP and magnesium occupy the ATP binding site in the active site, and it is empty in ATP binding site of human PRPS1 (2HCR). There is a SO_4_^2-^ and a PO_4_^3-^ in R5P binding site of human PRPS1 (2HCR) and human PRPS2, respectively.

Some crystal structures of hRPPS1 and its mutants had been previously solved. We compared the crystal structures of wild-type hPRPS1 (2HCR) and our cryo-EM structure of hPRPS2. We labeled the amino acids with differences in the matched monomer of hPRPS2 and hPRPS1 (2HCR). All these amino acids are located at the same position (**Figure 4B**).

The structure of hPRPS2 is very similar to that of hPRPS1 (2HCR). In the structure of hPRPS1 (2HCR), SO_4_^2-^ occupies the β-phosphate of ADP, and another SO_4_^2-^ binds to another regulator site (**Figure 4C**). From our structure, ADP can also bind to the allosteric site, and the binding of ADP will hinder the phosphate binding of allosteric sites. The active sites of hPRPS2 and hPRPS1 change little, and both phosphate and SO_4_^2-^ can bind to the R5P binding site of the active site (**Figure 4D**).

### Comparison of hexamer interface in human and *E. coli* PRPS

According to sequence alignment, we found the amino acids involved in connection of two adjacent hexamer are highly conserved in hPRPS and *E. coli* PRPS (**Figure 5A**). The hPRPS2 filament is similar to *E. coli* PRPS type A (7XMU) and type A^ADP+AMP^ (7XMV) filament. They have the same amino acid residues connecting adjacent hexamers. The positions of amino acid residues E298, R301, R302, N305 and E307 in the structures of hPRPS2 and *E. coli* PRPS type A filament (7XMU) and type A^ADP+AMP^(7XMV) filament do not changed significantly (**Figure 5B** and **C**).

**Figure 5.**
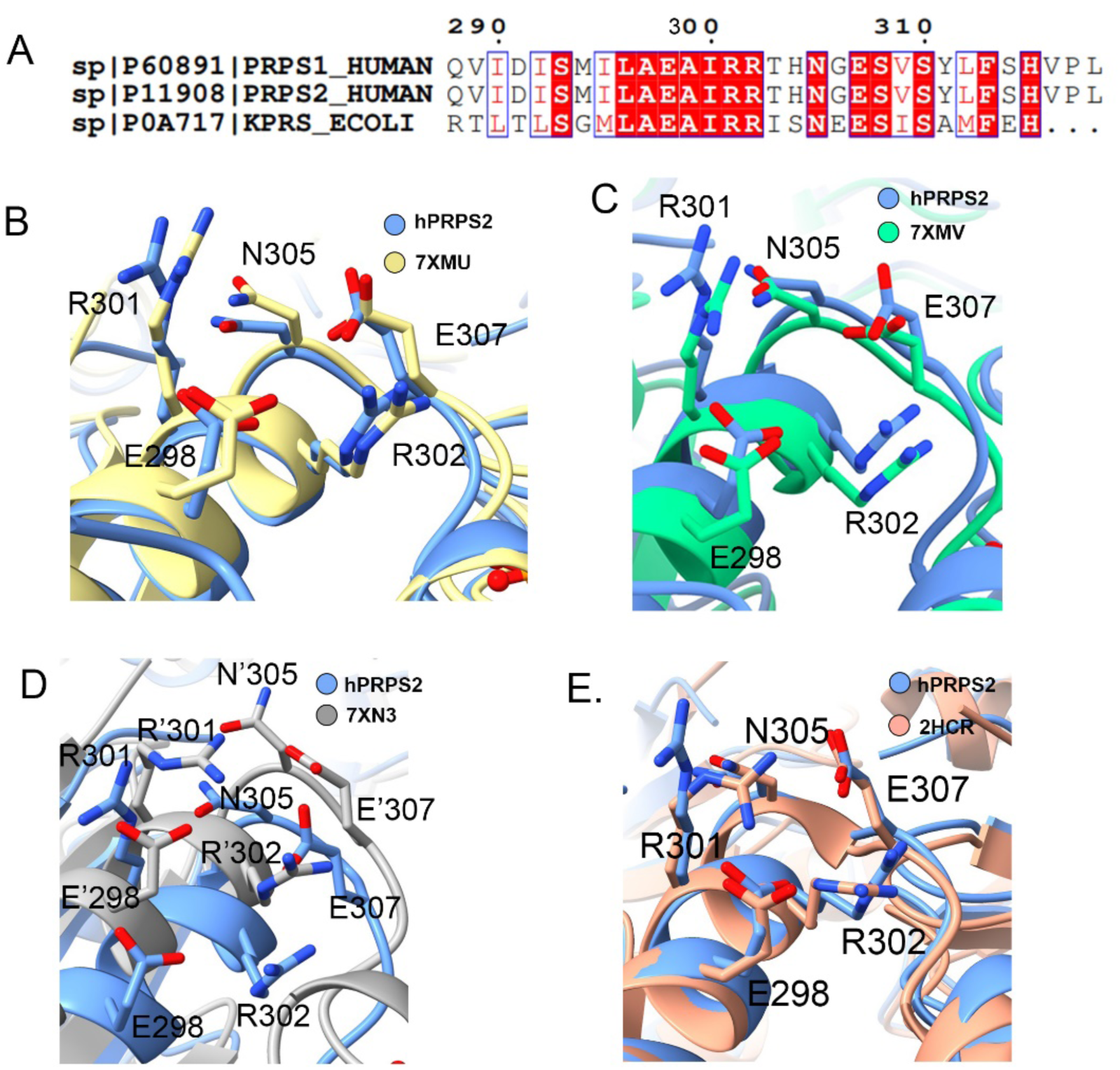
Structural comparison of interface amino acids in different organisms. (A) Sequence alignment of interface amino acids in different organisms. Some amino acids located at the interface of two adjacent hexamers in human PRPS2 filament are conserved. And the conserved amino acids are in red. (B) Structural comparison of interface amino acids between hPRPS2 (in blue) and *E. coli* type A filament PRPS (7XMU) (in yellow). Human PRPS2 filement is same to *E. coli* PRPS type A filament, and the amino acids involved in the interconnection of hexamers are consrved. (C) Structural comparison of interface amino acids between human PRPS2 (in blue) and *E. coli* type A^ADP+AMP^ filament PRPS (7XMV) (in green). The position of the amino acids involved in the interconnection of hexamers are highly similar. (D) Structural comparison of interface amino acids between human PRPS2 (in blue) and *E. coli* type B filament PRPS (7XN3) (in gray). Compared with *E. coli* type B filament PRPS (7XN3), the position of the amino acids involved in the interconnection of hexamers shifted. Amino acids in *E. coli* type B filament PRPS (7XN3) are labeled with ‘. (E) Structural comparison of interface amino acids between hPRPS2 and hPRPS (2HCR). There some amino acids involved in the interconnection of hexamers are not conserved in Bacillus subtilis PRPS. The amino acids in hPRPS (1DKU) are labeled with ‘.

*E.coli* PRPS type B filament has another interface between adjacent hexamers. There is a greater difference between hPRPS2 and *E. coli* type B filament (7XN3), and the amino acids involved in the formation of hPRPS2 filament have a great displacement relative to *E. coli* type B filament (**Figure 5D**). We also compared the crystal structure of hRPPS1 (2HCR) and hRPPS2, which have very similar structures at the interface between two adjacent hexamers of the filament (**Figure 5E**).

### Long and short forms of hPRPS2

There are two isoforms of hPRPS2: hPRPS2-long and hPRPS2-short. After the 102nd amino acid residue, hPRPS2-long contains three more amino acid residues than hPRPS2-short. In addition, among the compared organisms, only *E. coli* PRPS has an Ala after S103 in the RF loop (**Figure 6A**). This study solved the short isoform PRPS2 structure. When comparing the structure of hPRPS2 with that of *E. coli* type A (7XMU) and type A^ADP+AMP^ (7XMV) filaments, the regulatory flexible loop (RF loop Y94-T109) of *E. coli* clashes with ADP in human PRPS2 allosteric site 1. The RF loop conformation of hPRPS2 (Y94-S108) is very similar to that of *E. coli* PRPS including loops Y94-D101 and R104-S108 (Y94-V101 and R105-T109 in *E. coli*). The loop K102-S103 (R102-S103-A104 in E. coli) of hPRPS2 is shorter than that in *E. coli* and deviates from the *E. coli* loop by about 4.7Å (**Figure 6B** and **C**). The additional three amino acid residues of hPRPS2-long are located in the RF loop in the range of K102-S103, which results in a longer RF loop than *E. coli* PRPS. We speculate that the longer RF loop of the hPRPS2-long isoform may hinder the binding of ADP to allosteric site 1.

**Figure 6.**
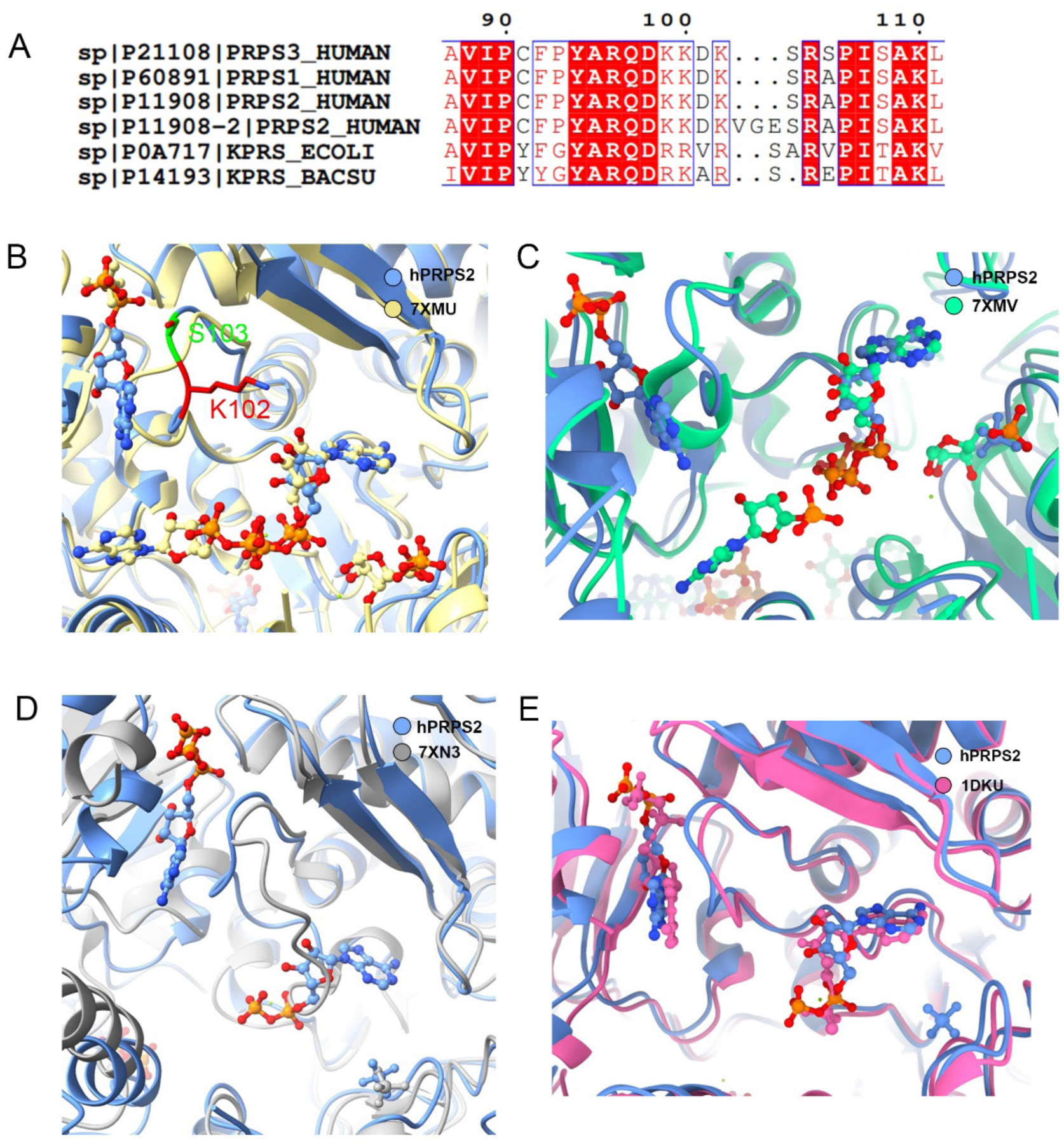
Ligands comparison of PRPS in different organisms. (A) The sequence alignment of RF loop of hPRPS1-3, *E. coli* PRPS and *Bacillus subtilis* PRPS. Structural comparison between human PRPS2 with *Escherichia coli* PRPS in type A filament (B: 7XMU), *Escherichia coli* PRPS in type A^ADP+AMP^ filament (C: 7XMV), *Escherichia coli* PRPS in type B filament (D: 7XN3), and *Bacillus subtilis* PRPS (E: 1DKU). In (B) and (C), ADP in the allosteric site 1 was clashed with the RF loop in *E. coli* PRPS. The difference of the RF loop in hPRPS2 short isoform is indicated (K102 and S103) in (B). In (D), ADP bound to the ATP active site of hPRPS2. And unlike the *E. coli* type B filament, the RF loop of hPRPS2 was not occupied the ATP site. In (E), the RF loop of human PRPS2 was slightly different from the *Bacillus subtilis* PRPS.

When comparing the structure of hPRPS2 with that of *E. coli* type B filament (7XN3), the RF loop of hPRPS2 does not occupy ATP sites as *E. coli* type B filament do (**Figure 6D**). This indicates that ADP binding at allosteric site 1 does not prevent ATP from entering its active site. While comparing the structure of hPRPS2 with that of *Bacillus subtilis* PRPS (1DKU), the RF loops between hPRPS2 and *Bacillus subtilis* PRPS are slightly different (**Figure 6E**).

## DISCUSSION

PRPS isolated from rat liver tissues revealed a heterogenous protein complex with a molecular weight greater than 1000 kDa (Liu, 2016)(Kita et al., 1989). A similar phenomenon was observed in human PRPS1/2 isolated from tissue sources (Becker et al., 1977; Foxi and Kelley, 1971). Furthermore, the recombinant human PRPS1/2 purified from *E. coli* can also be spontaneously assembled into large complexes larger than 1000 kDa in vitro (Nosal *et al*., 1993). Although these complexes are not necessarily filaments depicted in this study, collective evidence suggests that class I PRPS is regulated by the assembly of large complexes in bacteria and mammals.

Through large-scale screenings, dozens of metabolic enzymes including CTP synthetase (CTPS), inosine monophosphate dehydrogenase (IMPDH), asparagine synthetase (ASNs), phosphofructokinase (PFK) show the capability of forming filamentous structure termed cytoophidia. The cytoohidia structure has discovered to provide a common and conserved regulatory mechanism for many metabolic enzymes from archaea to prokaryotes and eukaryotes (Liu, 2016; Park and Horton, 2019; Zhou et al., 2020). As the rate limiting enzyme of purine and pyrimidine in organisms, PRPS was also found to form cytoophidia in a variety of eukaryotes including yeast, zebrafish and human (Begovich *et al*., 2020; Noree *et al*., 2019).

Taking advantage of cryo-EM and laser-scanning confocal microscopy, we have recently found that PRPS in E. coli forms cytoophidia as well. E. coli PRPS forms two types of filament in vitro and in vivo (Hu *et al*., 2022). The *E. coli* PRPS type A filament attenuates the allosteric inhibition by ADP, while the *E. coli* PRPS type B filament may influence the binding of ATP. hPRPS2 filament is similar to *E. coli* PRPS type A filament. While mutant R302A in hPRPS2 almost lost activity, mutant R302A in E. coli can function normally without allosteric regulator ADP exist. So there are some difference between the functions of hPRPS2 and E. coli filament.

Long isoform hPRPS2 has three more amino acids than short isoform after the 102nd amino acid, which are located at the RF loop. The longer RF loop may hinder ADP binding to allosteric site, which may be the reason hPRPS2 is insensitive to ADP/GDP inhibition. From our model, though ADP bind to the allosteric site, the RF loop do not occupy the ATP binding site of the active site. The ADP binding of allosteric site do not prevent ATP from entering the active site. The mechanism of ADP inhibition needs further exploration and research.

The structure comparison of hPRPS1 and hPRPS2 shows that they have same amino acids at the filament interface. Previous study found rat PRPS1 and PRPS2 as well as the two so-called PAP-39 and PAP-41 peptides can form big complex in rat liver (Kita *et al*., 1989). During the preparation of this manuscript, a preprint by Kollman and coworkers reported that hPRPS1 can form filaments (Hvorecny et al., 2022). We speculate that hPRPS1 and hPRPS2 may exist in the same filament. In the future, it would be interesting to solve the structure of the hPRPS1/hPRPS2 hybrid filaments.

## MATERIALS AND METHODS

### Human PRPS2 protein purification

Full-length wild-type or mutant human PRPS2 sequences with a C-terminal 6×His-tag were cloned into a modified pRSFDuet vector and expressed in *E. coli* Transetta (DE3) cells. After induction with 0.1 mM IPTG at the OD600 range of 0.5∼0.8, the cells were cultured at 37 °C for 4 hours and pelleted by centrifugation at 4,000 r.p.m. for 10 minutes. The harvested cells were sonicated under the ice in lysis buffer (50 mM Tris-HCl pH 8.0, 500 mM NaCl, 10% glycerol, 20 mM imidazole, 1 mM PMSF, 5 mM β-mercaptoethanol, 5 mM benzamidine, 2 μg/ml leupeptin and 2 μg/ml pepstatin). After ultrasonication, the cell lysate was then centrifuged (15,000 r.p.m.) at 4 °C for 45 minutes. The supernatant was collected and incubated with equilibrated Ni-NTA agarose beads (Qiagen) for 1 hour. and purified by Ni-NTA agarose beads (Qiagen). Lysis buffer with 50 mM imidazole was used to wash the column. And target proteins were eluted with lysis buffer with 250 mM imidazole. Further purification was performed in column buffer (25 mM Tris HCl pH 8.0 and 150 mM NaCl) using HiLoad Superdex 200 gel-filtration chromatography (GE Healthcare). The peak fractions were collected, concentrated, and stored in small aliquots at −80 °C. All the experiments were performed at 4 °C.

### Negative staining

Wild-type or mutation hPRPS2 proteins were mixed with different substrate conditions. In brief, purified hPRPS2 protein (1 μM) was dissolved in Tris-HCl buffer (25 mM Tris-HCl, 150 mM NaCl, 10 mM MgCl_2_), and 2 mM ligands (ATP, ADP, AMP, R5P, or PRPP) was added to the solution. After incubation at 37 °C for 30 min, the prepared protein samples were applied to glow-discharged carbon-coated EM grids (400 mesh, EMCN), and stained with 1% uranyl acetate. Negative-stain EM grids were photographed on a Tecnai Spirit G21 microscope (FEI).

### Cryo-EM grid preparation and data collection

6 μM hPRPS2 protein was dissolved in a buffer containing 25 mM Tris HCl pH 7.5, 150 mM NaCl, 2 mM ADP, and 10 mM MgCl_2_ to generate filaments. The samples were incubated on ice for 15 min and then loaded on H_2_/O_2_ glow-discharged amorphous alloy film (CryoMatrix M024-Au300-R12/13). Then Grids were immediately blotted for 3.0 s with blot force of -1 and plunge-frozen in liquid ethane cooled by liquid nitrogen using Vitrobot (Thermo Fisher Scientific) at 4 °C and with 100% humidity.

Movies were recorded on Titan Krios G3 (FEI) equipped with a K3 Summit direct electron detector (Gatan), operating in counting super-resolution mode at 300 kV. Each movie stack was acquired in a total dose of 60 *e*^−^Å^−2^, subdivided into 50 frames at 4 s exposure. Automated data acquisition was performed with SerialEM(Mastronarde, 2005) at a nominal magnification of 22,500× and a calibrated pixel size of 1.06 Å, with defocus ranging from 1.0 to 2.5 μm.

### Image processing and 3D reconstruction

All image processing steps were performed using Relion3.1-beta(Zivanov et al., 2018). MotionCor2 (Zheng et al., 2017) and CTFFIND4(Rohou and Grigorieff, 2015) were used to pre-process the image by RELION GUI. And the CTF (contrast transfer function) parameter was estimated by CTFFIND4. 681672 particles were auto-picked from 2403 micrographs. After 2D classification, 458154 particles were selected for 3D classification. After 3D classification using C1 and D3 symmetry, a total of 140303 particles of the best category were selected for 3D auto-refinement, and each particle was subjected to CTF refinement and Bayesian polishing. we get an initial 3.3 Å density map including three layers of hPRPS2 hexamer. A final 3.08 Å map was sharpened by post-process using a tight mask for the central hexamer with a B-factor of 45 Å^2^.

### Model building and refinement

The structure of hPRP2 from AlphaFold was applied for the initial model. The hexamer models were generated and then docked into the corresponding electron density map by Chimera v.1.14. Coot(Emsley and Cowtan, 2004)was used for iterative manual adjustment and rebuilding. The final atomic model was evaluated using MolProbity(Williams et al., 2018). The map reconstruction and model refinement statistics are listed in Supplementary Table 1. All figures were generated using UCSF Chimera(Pettersen et al., 2004) and ChimeraX(Goddard et al., 2018).

### PRPS activity assay

The activity of hPRPS2 was measured by coupled continuous spectrophotometry using SpectraMax i3. The production PRPP of PRPS can be determined by a coupled reaction (OA + PRPP → OMP + PPi) of *E. coli* orotate phosphoribosyltransferase (OPRT, EC 2.4.2.10). The amount of PRPP generated in the reaction was determined by the reduction of orotate (OA) in the mixture. The concentration of OA was measured by absorbance at 295 nm for 300 sec at 37 °C(Krungkrai et al., 2005). Reaction mixture (100 μl) contains 0.5 μM PRPS, 1 mM OPRT, 1 mM OA, 10 mM MgCl_2_, 250 mM NaCl, 30 mM Na_2_HPO_4_, 1 mM R5P or ATP at concentrations as described in each experiment. ATP or R5P was least added into the mixture to initiate the reaction. All measurements were performed in triplicate.

## ACKNOWLEGEMENTS

The EM data were collected at the ShanghaiTech Cryo-EM Imaging Facility. We thank the Molecular and Cell Biology Core Facility (MCBCF) at the School of Life Science and Technology, ShanghaiTech University for providing technical support. This work was supported by grants from Ministry of Science and Technology of China (No. 2021YFA0804700), National Natural Science Foundation of China (No. 31771490), Shanghai Science and Technology Commission (No. 20JC1410500) for grants to J.L.L.

## SUPPLEMENTARY FIGURES S1-S4 AND FIGURE LEGENDS

**Supplementary Figure S1.**
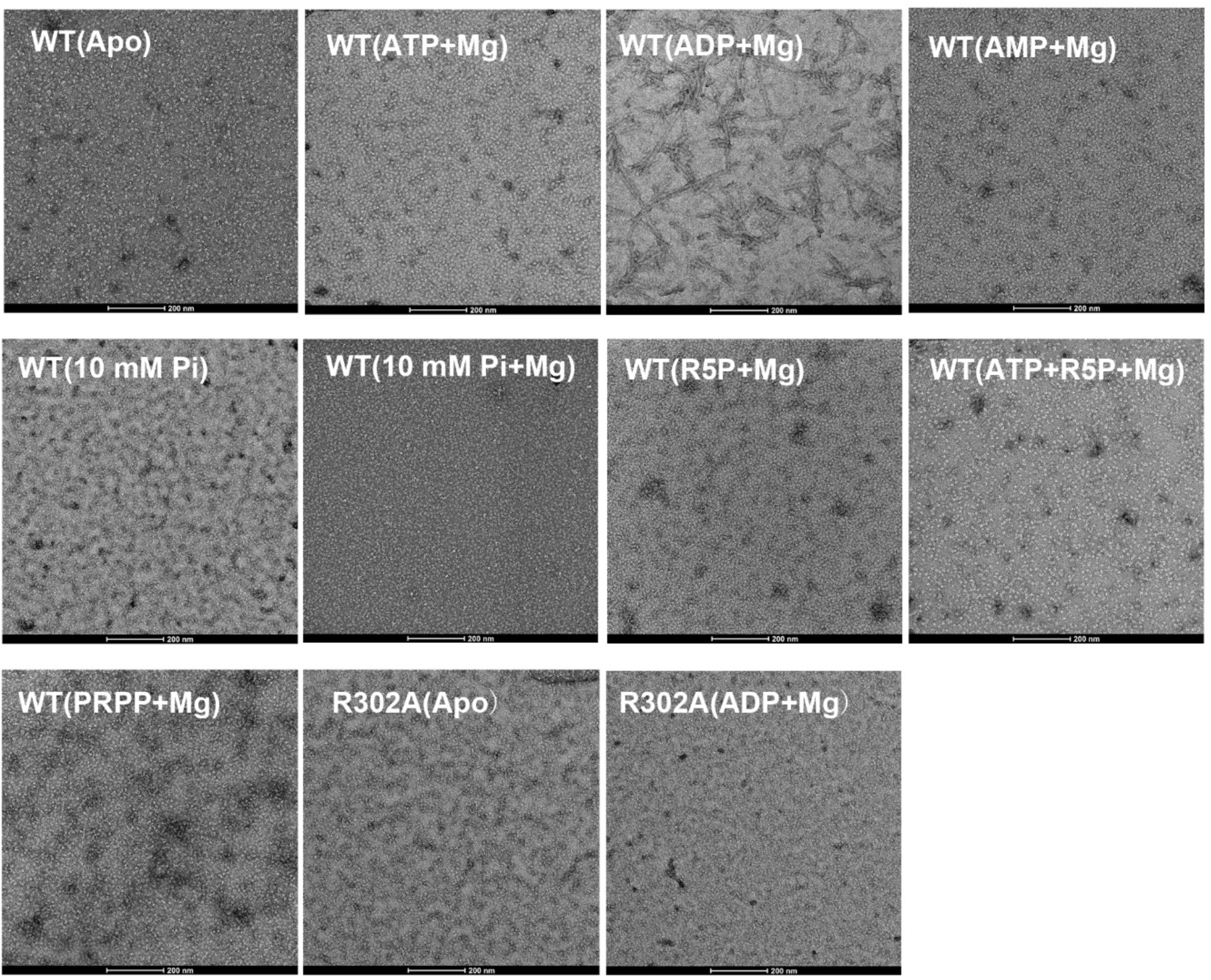
Human PRPS2 is assembled into filaments in vitro. Negative staining electron microscopic images of purified human PRPS2(1 μM) incubated in various conditions. The concentrations of nucleotides, phosphate ions (Pi) and Mg^2+^ are 2 mM, 30mM and 10 mM, respectively.

**Supplementary Figure S2.**
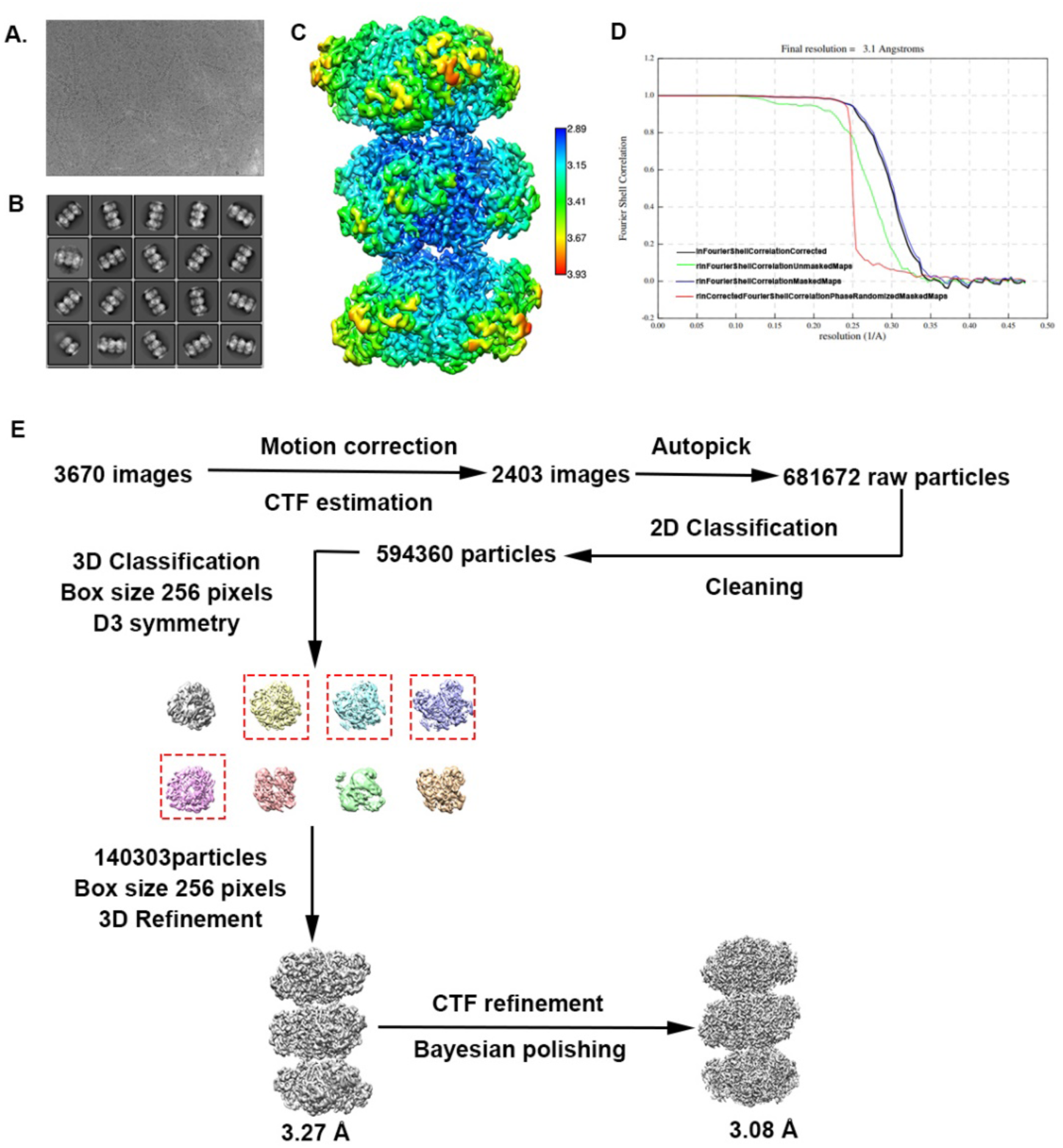
Cryo-EM data processing of human PRPS2 filament. (A) Representative Cryo-EM image of human PRPS2 filament. (B) Representative 2D averages of human PRPS2 filament in different views. (C) Local resolution of the type B filament final density map. (D) FSC curves of central hexamer in human PRPS2 filament density map (dash line shows FSC = 0.143). The final average resolution of hexamer is estimated to be 3.1 Å. (E) Flow chart of human PRPS2 filament image processing.

**Supplementary Figure S3.**
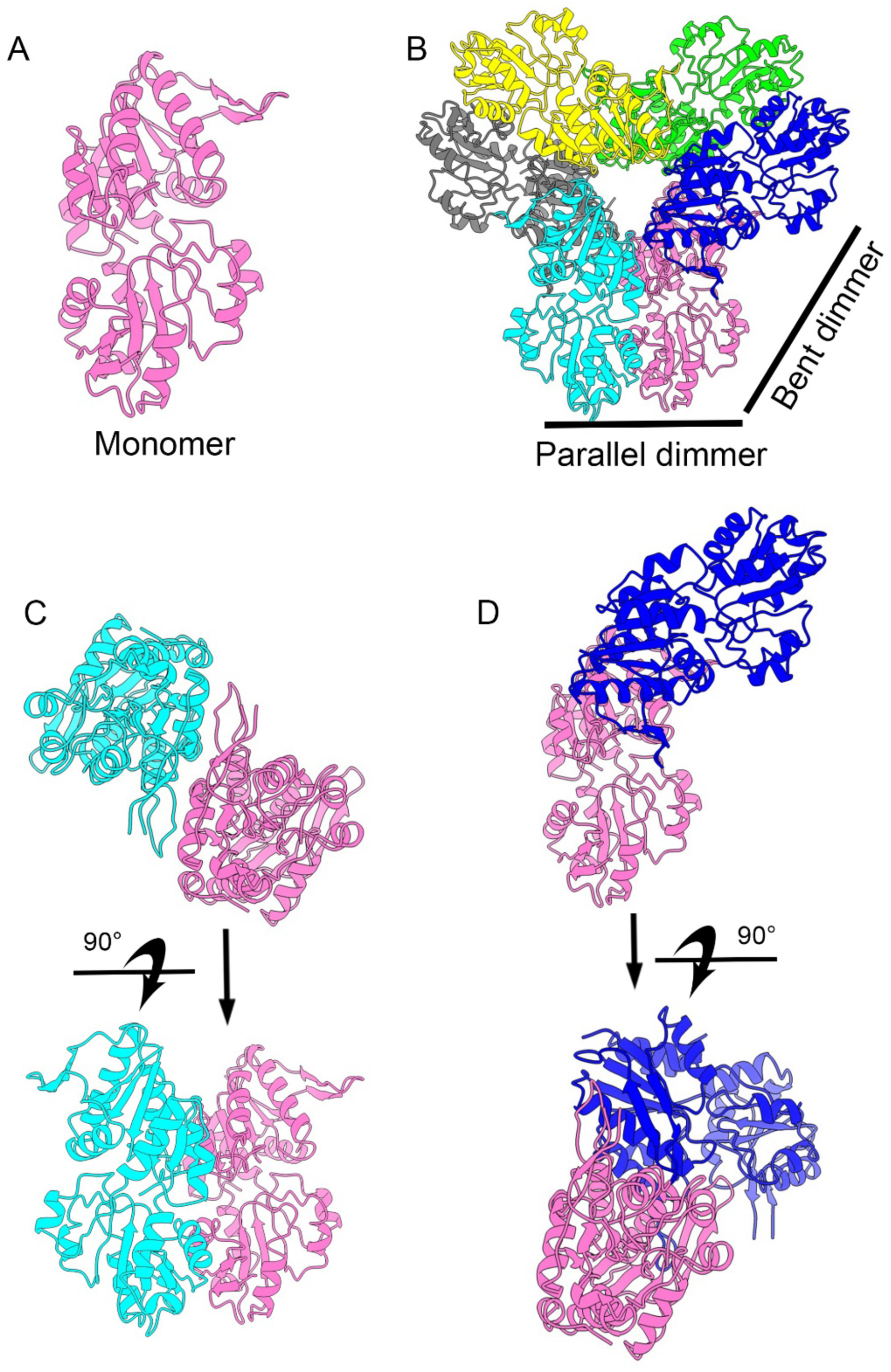
The structure of human PRPS2. (A) Monomer of human PRPS2. (B) The hexamer of human PRPS2. Each chain is in different color. (C) The parallel dimmer of human PRPS2. Each chain is in different color. (D) The bent dimmer of human PRPS2. Each chain is in different color.

**Supplementary Figure S4.**
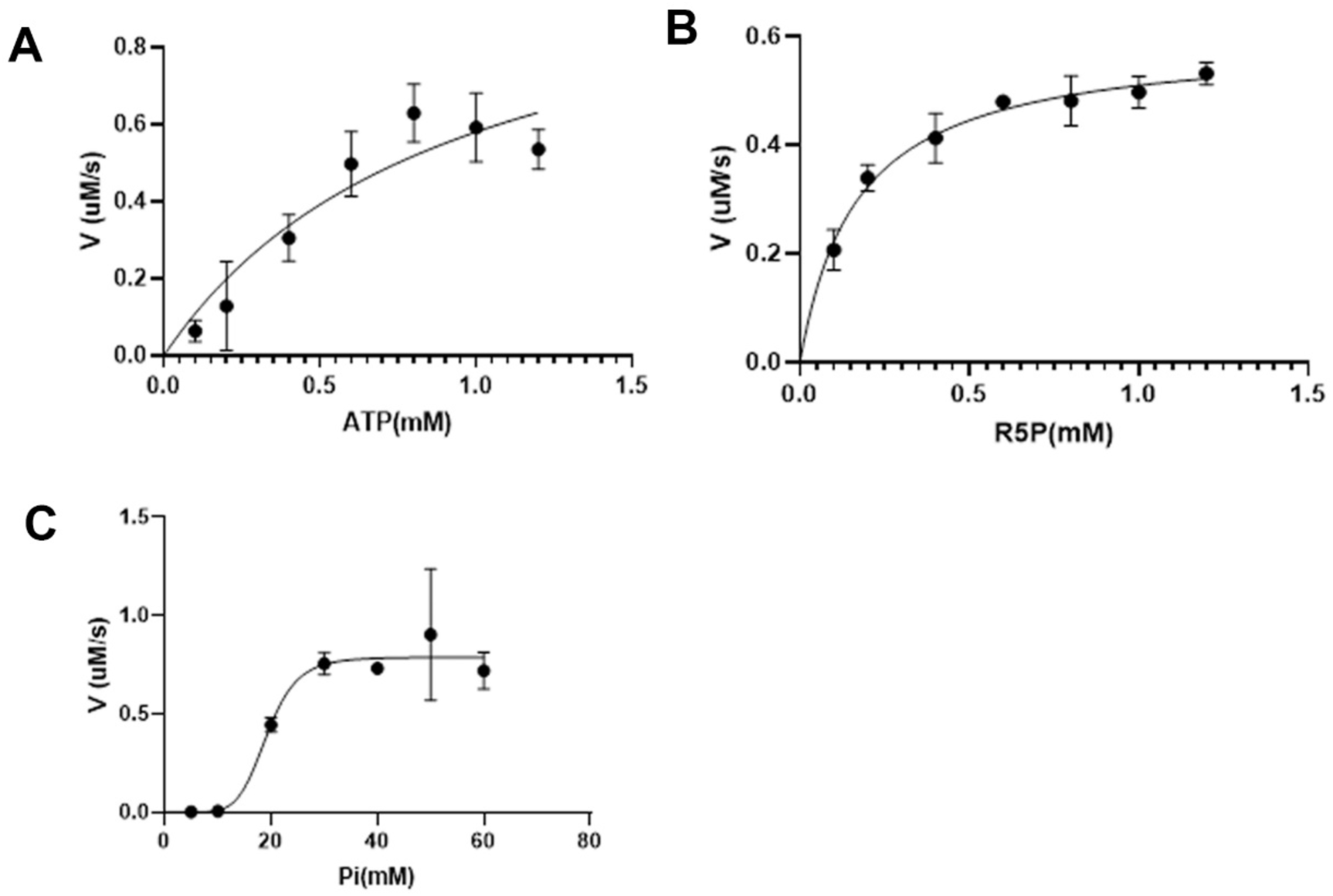
Catalytic activity of human PRS2 with different concentrations of ligands. Graphs show the catalytic activity of wild-type human PRPS2 in the presence of different amounts of ATP (A), R5P (B), and phosphate ion (C). All tests are repeated three times.

